# Genomic diversity across *Candida auris* clinical isolates shapes rapid development of antifungal resistance *in vitro* and *in vivo*

**DOI:** 10.1101/2022.03.25.485898

**Authors:** Laura S. Burrack, Robert T. Todd, Natthapon Soisangwan, Nathan P. Wiederhold, Anna Selmecki

## Abstract

Antifungal drug resistance and tolerance poses a serious threat to global public health. In the human fungal pathogen, *Candida auris*, resistance to triazole, polyene, and echinocandin antifungals is rising, resulting in multidrug resistant isolates. Here, we use genome analysis and *in vitro* evolution of seventeen new clinical isolates of *C. auris* from clades I and IV to determine how quickly resistance mutations arise, the stability of resistance in the absence of drug, and the impact of genetic background on evolutionary trajectories. We evolved each isolate in the absence of drug as well as in low and high concentrations of fluconazole. In just three passages, we observed genomic and phenotypic changes including karyotype alterations, aneuploidy, acquisition of point mutations, and increases in MIC values within the populations. Fluconazole resistance was stable in the absence of drug, indicating little to no fitness cost associated with resistance. Importantly, two isolates substantially increased fluconazole resistance to ≥256µg/ml fluconazole. Multiple evolutionary pathways and mechanisms to increase fluconazole resistance occurred simultaneously within the same population. Strikingly, the sub-telomeric regions of *C. auris* were highly dynamic as deletion of multiple genes near the sub-telomeres occurred during the three passages in several populations. Finally, we discovered a mutator phenotype in a clinical isolate of *C. auris*. This isolate had elevated mutation rates compared to other isolates and acquired substantial resistance during evolution *in vitro* and *in vivo* supporting that the genetic background of clinical isolates can have a significant effect on evolutionary potential.

**Importance:** Drug resistant *Candida auris* infections are recognized by the CDC as an urgent threat. Here, we obtained and characterized a set of clinical isolates of *C. auris* including multiple isolates from the same patient. To understand how drug resistance arises, we evolved these isolates and found that resistance to fluconazole, the most commonly prescribed antifungal, can occur rapidly and that there are multiple pathways to resistance. During our experiment, resistance was gained, but it was not lost, even in the absence of drug. We also found that some *C. auris* isolates have higher mutation rates than others and are primed to acquire antifungal resistance mutations. Furthermore, we found that multidrug resistance can evolve within a single patient. Overall, our results highlight the high stability and high rates of acquisition of antifungal resistance of *C. auris* that allow evolution of pan-resistant, transmissible isolates in the clinic.

## Introduction

*Candida auris* has spread rapidly since it was first identified as a fungal species in 2009 in Japan [1]. *C. auris* infections have been reported in at least 47 countries, although given challenges in distinguishing *C. auris* from other species, infections are likely even more widespread [2]. Genomic analysis indicates that *C. auris* isolates can be classified into five major geographically distinct clades with clades I, III and IV most commonly causing outbreaks [3–5]. Major factors contributing to the rapid spread of *C. auris* are high rates of transmission from patient-to-patient in healthcare settings, extended survival on fomites, and rapidly acquired antifungal drug resistance [6, 7]. Acquired resistance to triazoles, including fluconazole, as well as polyenes is common, and resistance to echinocandin antifungals is increasing, resulting in multidrug resistant isolates [8]. Pan-resistant isolates of *C. auris* have recently arisen and spread from one patient to another during outbreaks in multiple healthcare facilities in the United States [9]. Therefore, understanding how *C. auris* becomes resistant to antifungals is critical.

Azole resistance in *Candida* species most often occurs due to mutations within the ergosterol biosynthesis pathway. Mutations in *ERG11,* the gene that encodes a sterol-demethylase and is the target of triazoles, are commonly identified in drug resistant clinical isolates in addition to mutations that activate drug export pathways. In *C. auris* isolates, fluconazole resistance has been correlated with three amino acid substitutions in *ERG11*: V125A/F126L, Y132F, and K143R [4]. These mutations have been shown to increase fluconazole minimum inhibitory concentrations (MICs) by ∼16-fold [10]; however, mutations in *ERG11* alone cannot explain the very high levels of resistance (MICs >256µg/ml fluconazole) observed in many *C. auris* isolates. Activating mutations in the transcriptional regulator of drug efflux pumps, *TAC1B*, have also been shown to be important for high levels of fluconazole resistance [11], and deletion of *TAC1B* abrogates resistance [12]. However, there are still many open questions regarding the rates and stability of acquired drug resistance and the contribution of genetic diversity between clinical isolates to the evolutionary rates and trajectories of acquired drug resistance.

*In vitro* evolution experiments using single *C. auris* isolates have provided insight into the rates of evolution and types of mutations acquired during drug exposure. Evolution of the clade I reference genome strain AR0387/B8441 in increasing concentrations of fluconazole followed by candidate gene sequencing highlighted the importance of *TAC1B* as a key regulator of resistance [11]. Passaging of a different fluconazole susceptible clade I isolate in increasing concentrations of fluconazole identified aneuploidy of Chromosome 5 along with missense mutations in three genes, including *TAC1B*. Chromosome 5 contains *TAC1B* as well as several other possible fluconazole resistance genes. Increased copy number elevates transcription of these genes [13] similar to aneuploidy-based fluconazole resistance in *Candida albicans* [14, 15]. A drug sensitive clade II isolate evolved resistance in <21 days after passaging in increasing concentrations of amphotericin B, fluconazole, or caspofungin. Furthermore, passaging in amphotericin B or combinations of fluconazole and caspofungin resulted in multidrug resistance in the clade II isolates [16]. Resistant isolates acquired missense mutations and/or aneuploidies. For example, the isolates passaged in amphotericin B acquired missense mutations in *ERG3, ERG11,* and *MEC3,* genes associated with increased MICs for both amphotericin B and fluconazole, while the multidrug resistant isolate passaged in fluconazole and then caspofungin had mutations in *TAC1B, FKS1* (the target of caspofungin), and a segmental duplication of chromosome 1 containing *ERG11* [16].

However, these studies have only used a single isolate such that the spectrum of mutations identified has been limited and the contributions of genetic background and starting progenitor fitness have not been determined. The spectrum of mutations known to increase fluconazole resistance is much broader in the better studied organism *C. albicans* than in *C. auris*. It is not known if this is because of differences in genome structure or other factors. *C. albicans* is typically a heterozygous diploid and therefore may be more likely to develop a wider range of aneuploidies, loss of heterozygosity events, and copy number variations based on repeat-based genome rearrangements than *C. auris*, which has a haploid genome [15,17–19]. However, recessive single nucleotide variations (SNVs) are initially buffered by the other homologous chromosome in *C. albicans*. *C. auris* is haploid so SNVs may have a larger impact on the rate/dynamics of adaptation, as has been seen in direct comparisons of evolutionary pathways in diploid and haploid strains of *Saccharomyces cerevisiae* [20, 21]. *Candida glabrata* is also haploid and prone to developing multidrug resistance. Recent experimental evolution studies with *C. glabrata* showed that a relatively small number of genes are mutated during the acquisition of drug resistance, but that the range of SNVs within these genes is high [22]. Understanding the spectrum of aneuploidies, genes, and SNVs associated with resistance in *C. auris* will help provide insight into clinically relevant markers of resistance.

Antifungal drug tolerance is defined as a subset of cells that can grow slowly in drug concentrations above the MIC, and is correlated with treatment failure in the clinic [23, 24]. In *C. auris* tolerance to azoles requires the essential molecular chaperone *HSP90* [25], is enhanced as mother cells age, and is associated with gene duplication and overexpression of *ERG11* and *CDR1*, encoding an ABC transporter [26]. To date, *in vitro* evolution experiments in *C. auris* have focused on acquisition of resistance through MIC measurements [13, 16], whereas the range of tolerance observed in clinical isolates and changes in tolerance over time have not been studied. Understanding the contribution of genetic background to drug resistance and tolerance is particularly important because mutational and phenotypic outcomes depend on the starting genetic landscape of the organism [27, 28]. For example, different clinical isolates of *C. albicans* have heterogeneous drug responses and variable levels of genome stability in fluconazole [18,23,29].

Mutations and copy number variations can provide enhanced fitness in the presence of the antifungal drug, but these resistance mutations often incur a fitness cost in the absence of the drug [19,22,30]. Multiple studies in *C. albicans* have shown rapid loss of antifungal drug resistance mutations in the absence of selective pressure [30, 31]. However, little is known about the stability of drug resistance alleles in *C. auris,* and none of the *in vitro* evolution studies so far have evolved clinically drug resistant isolates in the absence of antifungals or in sub-inhibitory concentrations to measure the relative rates of loss of resistance compared to gain.

Here, we evolved a set of seventeen clinical isolates of *C. auris* from clades I and IV to determine how quickly resistance mutations arise, the prevalence of tolerance, the stability of resistance in the absence of drug, and the influence of genetic background on antifungal drug resistance. In the presence and absence of fluconazole, we observed genomic and phenotypic changes including karyotype alterations, aneuploidy, loss of sub-telomeric regions, acquisition of *de novo* point mutations, and increases in MIC values and tolerance within the populations. Strikingly, we observed little to no fitness cost associate with resistance as the clinical isolates with high starting MICs maintained fluconazole resistance in the absence of drug. We also found that the genetic background of clinical isolates dramatically impacts evolutionary dynamics. One clinical isolate had elevated mutation rates relative to other isolates and acquired substantial resistance during the evolution experiment. This is the first example of a mutator phenotype detected in clinical isolates of *C. auris*. In lineages from this isolate, multiple mechanisms to increase fluconazole resistance occurred simultaneously within the same population, including missense alleles in transcriptional regulators of azole resistance and acquisition of multiple aneuploidies. Retrospective analysis of clinical data identified that this mutator acquired multidrug resistance during infection of an individual patient. Overall, our results demonstrate the high potential for rapid evolvability of drug resistance in clinical isolates of *C. auris*.

## Results

### Genomic diversity among seventeen clinical isolates of *Candida auris*

To better understand how genomic diversity contributes to drug resistance and infection potential of *Candida auris*, we obtained seventeen *C. auris* isolates from the Fungus Testing Laboratory at the University of Texas Health Science Center. These isolates were isolated from diverse body sites (blood, urine, tissue samples, tracheal aspirate, and pleural fluid) and geographic locations (Table S1) [32, 33]. Available clinical information indicates that these 17 isolates were likely from 14 individual patients [33]. Using Illumina-based whole genome sequence (WGS), we determined that these isolates are members of *C. auris* clades I and IV (Figure 1A). Four previously characterized isolates [4], B8441, B11220, B11221, and B11245 were used as controls to identify clades I-IV (Figure 1A). We named the six clade I clinical isolate progenitors A-P to F-P and eleven clade IV clinical isolate progenitors G-P to Q-P. Among samples geographic location information, clade I samples were from New York, New Jersey and Massachusetts, while clade IV samples were from Illinois and Florida (Table S1). Isolates H-P, I-P and K-P are known to be from a single patient as are O-P and P-P.

**Figure 1:**
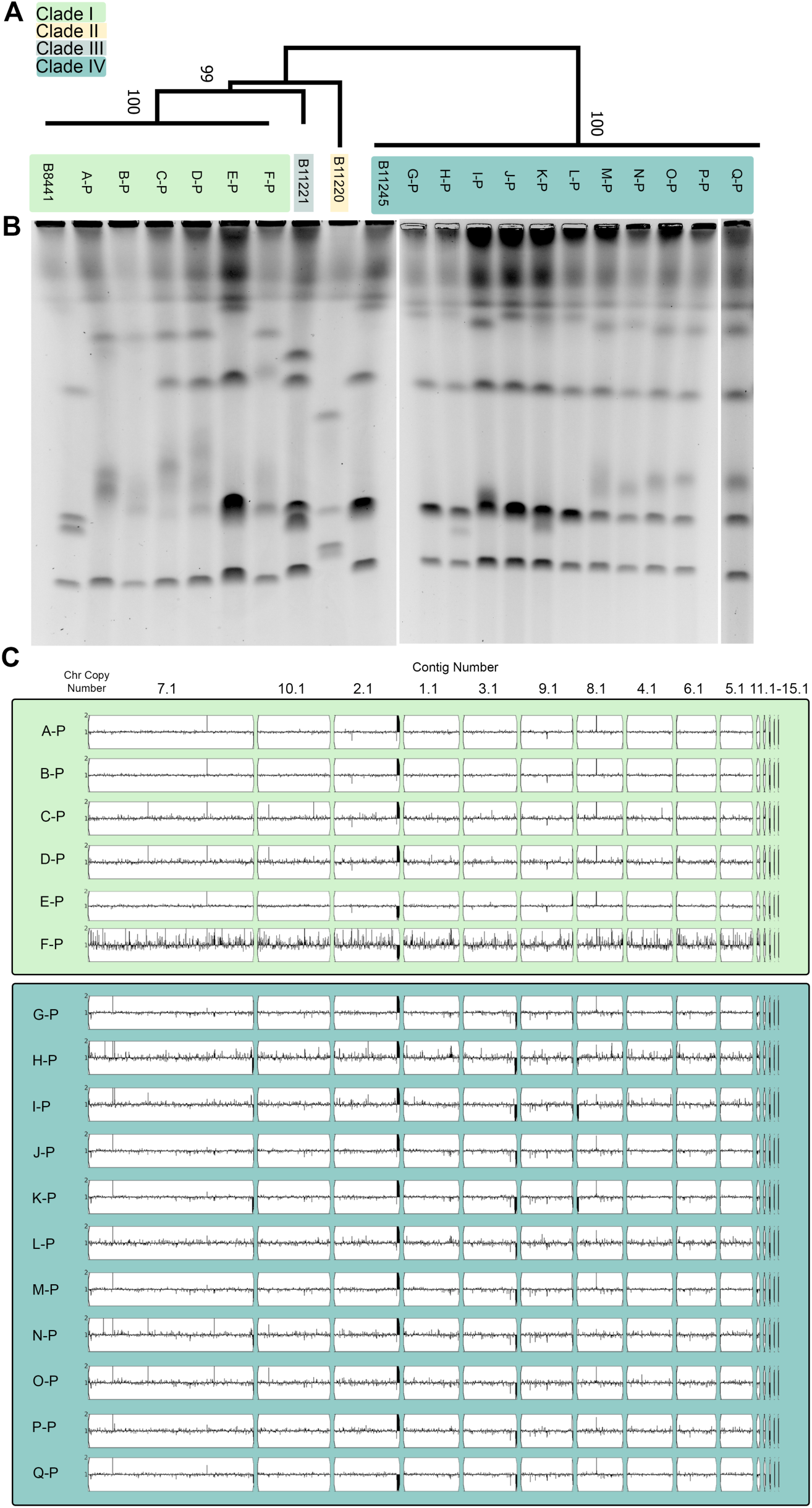
Characterization of the genomes of the seventeen new clinical isolates. (A) Phylogenomic analysis of the seventeen starting clinical isolates based on SNV data from Illumina whole-genome sequencing using the maximum likelihood method based on informative positions. Numbers along branches indicate bootstrap values calculated using 500 replicates. Clinical isolate strains A-P through F-P cluster to Clade I, while clinical isolates G-P through Q-P cluster to Clade IV. (B) CHEF karyotype gel stained with ethidium bromide of each clinical isolate as well as four previously characterized strains: B8441, B11221, B11220 and B11245 that represent clades I through IV, respectively. (C) Whole-genome sequence data plotted using YMAP. Read depth was converted to copy number (Y-axis) and was plotted as a function of chromosome position using the B8441 Clade I reference genome for each clinical isolate.

We next analyzed karyotypes using contour-clamped homogenous electric field (CHEF) electrophoresis. Overall, the karyotypes of isolates within a clade were more similar than karyotypes across clades, but we also observed high intra-clade karyotype variability. Strikingly, the karyotype of every clade I isolate differed from the other clade I clinical isolates, and four distinct karyotypes were observed among the eleven clade IV clinical isolates (Figure 1B). Interestingly, differences are apparent even among isolates from a single patient as I-P has a different karyotype from H-P and K-P. These results demonstrate chromosomal-level differences between clades of *C. auris* as well as among isolates from a single clade [5,34,35]. However, the mechanisms driving these chromosomal differences, such as copy number variations (CNVs), are not understood.

To identify potential CNVs, we mapped the Illumina WGS reads to the B8441 reference genome contigs using YMAP [36]. None of the clinical isolates had detectable whole chromosome aneuploidies, but we did observe CNVs in several clinical isolates, especially in sub-telomeric regions (Figure 1C). The CNV at the right edge of contig 2.1 aligns with the rDNA repeat region, and the internal CNVs on contigs 7.1, 4.1, 9.1 and 8.1 map to repetitive sequences indicating isolate-to-isolate variability in repeat copy number. For example, one of the amplifications visible on contigs 7.1 and 8.1 maps to an open reading frame (ORF) encoding a homolog of the *Zorro3* retrotransposase, demonstrating variable transposon copy numbers between isolates (Figure S1). Intriguingly, the sub-telomeric regions of several contigs have no copies remaining on the YMAPs (Figure 1C), so we analyzed read depth across these regions more closely [37]. We identified sub-telomeric deletions in the clinical isolates as regions greater than 500bp within 50kb of the end of the chromosome without reads mapped to the reference genome (Figure S2). Remarkably, every clinical isolate in our collection had at least one sub-telomeric deletion compared to the reference genome B8441 (Figure 1C, Table S1 and Table S2). Clade I clinical isolates had one or two sub-telomeric deletions totaling 4.6kb to 10.1kb, while clade IV clinical isolates had five or six sub-telomeric deletions totaling 39.5kb to 103.8kb (Table S1 and Table S2). These results indicate that *C. auris* sub-telomeric regions are highly dynamic and may account for some of the karyotype plasticity observed by CHEF (Figure 1B).

GO term analysis of the annotated gene functions of ORFs within the deleted sub-telomeric regions showed significant enrichment for transmembrane transporter activity (31.2% of genes, corrected p-value 1.45e-10). For example, isolate I-P was missing 21 ORFs including 2 predicted transmembrane iron transporters, 3 putative transmembrane glycerol transporters, and 2 putative transmembrane glucose transporters (Table S2). Within a clade, multiple clinical isolates shared sub-telomeric deletion patterns indicating that these isolates may be closely related. For example, five of the six clade I isolates shared the same deletion of 4.6kb at the end of contig 9.1. Four of these five had only this one deletion, but isolate A-P was also missing a portion of contig 7.1. The remaining clade I isolate (E-P) matched the reference sequences in contigs 9.1 and 7.1 but lacked a region at the beginning of contig 8.1 (Table S2). Consistent with this data, E-P had a distinct karyotype from the other clade I isolates despite overall similarity at the level of SNVs (Figure 1). These sub-telomeric deletions may result in phenotypic differences important for host adaptation and infection, such as iron import, between clades and between isolates within the same clade.

### Clinical isolates of *C. auris* have differing fluconazole susceptibility and tolerance profiles

We next used minimal inhibitory concentration (MIC_50_) values to quantify both resistance and tolerance of the clinical isolates. For some isolates, clinical data on resistance was collected at the Fungus Testing Lab, so we used this as a starting point to understand drug susceptibility. Based on these data, six of the seventeen isolates (A-P, B-P, C-P, D-P, H-P and K-P) were resistant to at least one antifungal drug class and three isolates (A-P, B-P and K-P) were resistant to two different classes of antifungal drugs using resistance breakpoints defined by the CDC [38] (Table S3). However, some isolates did not have clinical data for fluconazole. Additionally, the clinical tests do not include information on tolerance to fluconazole. Therefore, we measured fluconazole resistance (FLC^R^) and tolerance for all isolates. Five of the six clade I clinical isolates had a MIC_50_ of ≥256µg/ml fluconazole, while the sixth isolate had an MIC_50_ of 64µg/ml fluconazole (Table S1). In comparison, nine isolates from clade IV had MIC_50_ values ranging from 2µg/ml to 64µg/ml, with only a single FLC^R^ isolate K-P. We were unable to measure the MIC_50_ for isolate N-P at 24 hours because the isolate grows poorly *in vitro* and did not reach the culture density necessary for determining MIC_50_.

Antifungal drug tolerance is defined as the ability of cells to grow at concentrations of drug higher than the MIC_50_ and can be measured by supra-MIC growth (SMG) at 48 hours [23,24,39]. We measured SMG for all isolates with MIC_50_ values below 256µg/ml. Two of the reference isolates, B11220 and B11221, and one of the clinical isolates from this set, H-P had SMG values >0.6 indicating high levels of tolerance in these isolates. Furthermore, isolates E-P, K-P and Q-P had SMG values between 0.4-0.6. Therefore, tolerance occurs in *C. auris* and varies between clinical isolates similar to what has been observed in other *Candida* species [23, 29].

To identify potential drivers of antifungal drug resistance and tolerance, we performed comparative genomics to identify SNVs in the isolates relative to the reference isolate B8441, which is drug sensitive and has low tolerance. We first analyzed SNVs in *FKS1, ERG11, ERG3, TAC1B,* and *MRR1A* because of their previous connections to drug resistance in *C. auris*. *FKS1* encodes the echinocandin target 1,3-beta-D-glucan synthase and missense mutations have been previously linked to echinocandin resistance in *C. auris* [40–42]. Isolates H-P and K-P, the two isolates that were resistant to caspofungin, had *FKS1*^S639P^ missense mutations, while sensitive isolates did not have *FKS1* substitutions (Table S1 and Table S3). Loss of function of *ERG3*, which encodes a sterol Δ⁵,⁶-desaturase in the ergosterol biosynthesis pathway, has been linked to several types of drug resistance in other *Candida* species [22, 43]. In *C. auris*, S58T substitutions in *ERG3* have potentially been linked to amphotericin B resistance in a small set of clade IV clinical isolates [42]. We observed an *ERG3*^S58T^ variant in all the clade IV isolates, however we did not observe any correlation between this mutation and amphotericin B resistance. Strikingly, we observed a novel SNV in *ERG3* (T227I) in the H-P isolate which had high fluconazole tolerance with SMG values >0.6, suggesting that this allele may be correlated with tolerance. SNVs in *ERG11,* the target of fluconazole, and *TAC1B*, a regulator of drug export pumps, correlate with azole resistance in published datasets [4,11,16,44]. The five FLC^R^ isolates with MIC_50_ values ≥256µg/ml contained the mutations *ERG11*^K143R^ and *TAC1B*^A640V^ (Table S1). Both isolates (E-P and K-P), with MIC values in the 32-64µg/ml range had an *ERG11* missense mutation, *ERG11*^Y132F^, that has also been correlated with resistance (Table S1). We did not observe any *MRR1A* mutations associated with fluconazole resistance in our clade I or clade IV isolates.

To expand beyond the candidate gene approach to identify genetic variation that may explain higher levels of tolerance, we used comparative genomics of clade IV isolates with similar MICs, but varying degrees of tolerance. We compared two strains with high tolerance, H-P and Q-P, to a strain with low tolerance, I-P to identify moderate or high impact SNVs (e.g., missense and nonsense). There were 21 moderate and high impact SNVs in H-P that were not in I-P. In addition to the *ERG3* mutation discussed above, there were several other candidates that may affect transcriptional regulation or membrane composition based on annotated function (Table S4). There were 5 moderate and high impact SNVs in Q-P that were not in I-P. None of them were shared with H-P, the other high tolerance clinical isolate, suggesting that there are multiple mechanisms that can promote tolerance in *C. auris*. None of the identified SNVs in Q-P have been previously annotated as being involved in fluconazole responses, but two of the five have annotated functions related to mating in other fungi (Table S4). Together, these mutational analyses provide insight into mechanisms of pre-existing antifungal drug resistance and SNVs associate with tolerance within the clinical isolates.

### *In vitro* evolution of *Candida auris* in the presence and absence of fluconazole

To gain a better understanding of how genetic background and pre-existing antifungal drug resistance shape genome dynamics and evolution, we conducted a short-term *in vitro* evolution experiment using these clinical isolates as progenitors for approximately 30 generations in high concentrations of fluconazole (256µg/ml), sub-MIC concentrations of fluconazole (1µg/ml fluconazole), and in medium lacking antifungal drug (Figure 2). The same starting population was used for each of the environmental conditions. Serial dilution (1:1000) was conducted every 72 hrs for a total of three passages, and then the entire population of cells from each lineage were collected for MIC, ploidy, CHEF, and Illumina WGS analysis (Figure 2). We chose the 0µg/ml and 1µg/ml fluconazole conditions to gain insight into evolutionary pathways of *C. auris* in the absence of drug selection and in sub-MIC concentrations. Neither the 0µg/ml or 1µg/ml fluconazole condition resulted in dramatic changes in MIC (Table S1). Thirteen of the sixteen isolates had no change in MIC when evolved in the absence of drug. The remaining three isolates decreased their MICs by 2-fold (Table S1). No changes in MICs were observed for any of the isolates following passaging in the 1µg/ml fluconazole condition (Table S1). Therefore, resistance is maintained in sub-MIC levels of drug and in the absence of drug for at least 30 generations. In 256µg/ml fluconazole, the frequency of MIC change depended on starting MIC_50_ values. For progenitors with either low (≤4µg/ml) or very high (≥256µg/ml) MIC_50_, three passages in 256µg/ml fluconazole did not result in any change in MIC values. However, for the two progenitors (E and K) with MIC_50_ values of 64µg/ml fluconazole, passaging in 256µg/ml fluconazole increased the MIC_50_ in the evolved population to 128-256µg/ml fluconazole (K-256 and E-256) (Table S1 and Figure 3A).

**Figure 2:**
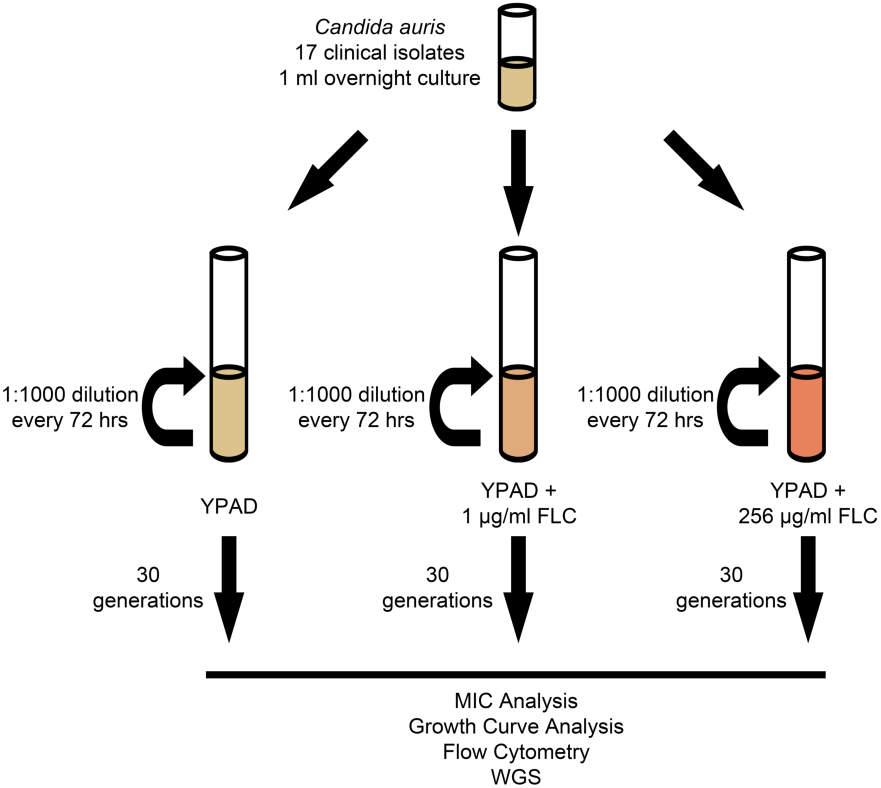
Schematic of *C. auris in vitro* evolution experiment. Seventeen *C. auris* clinical isolates were split into three independent lineages and cultured at three different concentrations of FLC (YPAD, YPAD + 1ug/ml FLC, & YPAD + 256ug/ml FLC) for a total of 51 evolution populations. Serial passage (1:1000 dilution) of each population occurred every 72 hrs for a total of 30 generations. After 30 generations, each of the 50 evolution populations were collected for further analysis (N-256 did not grow).

**Figure 3:**
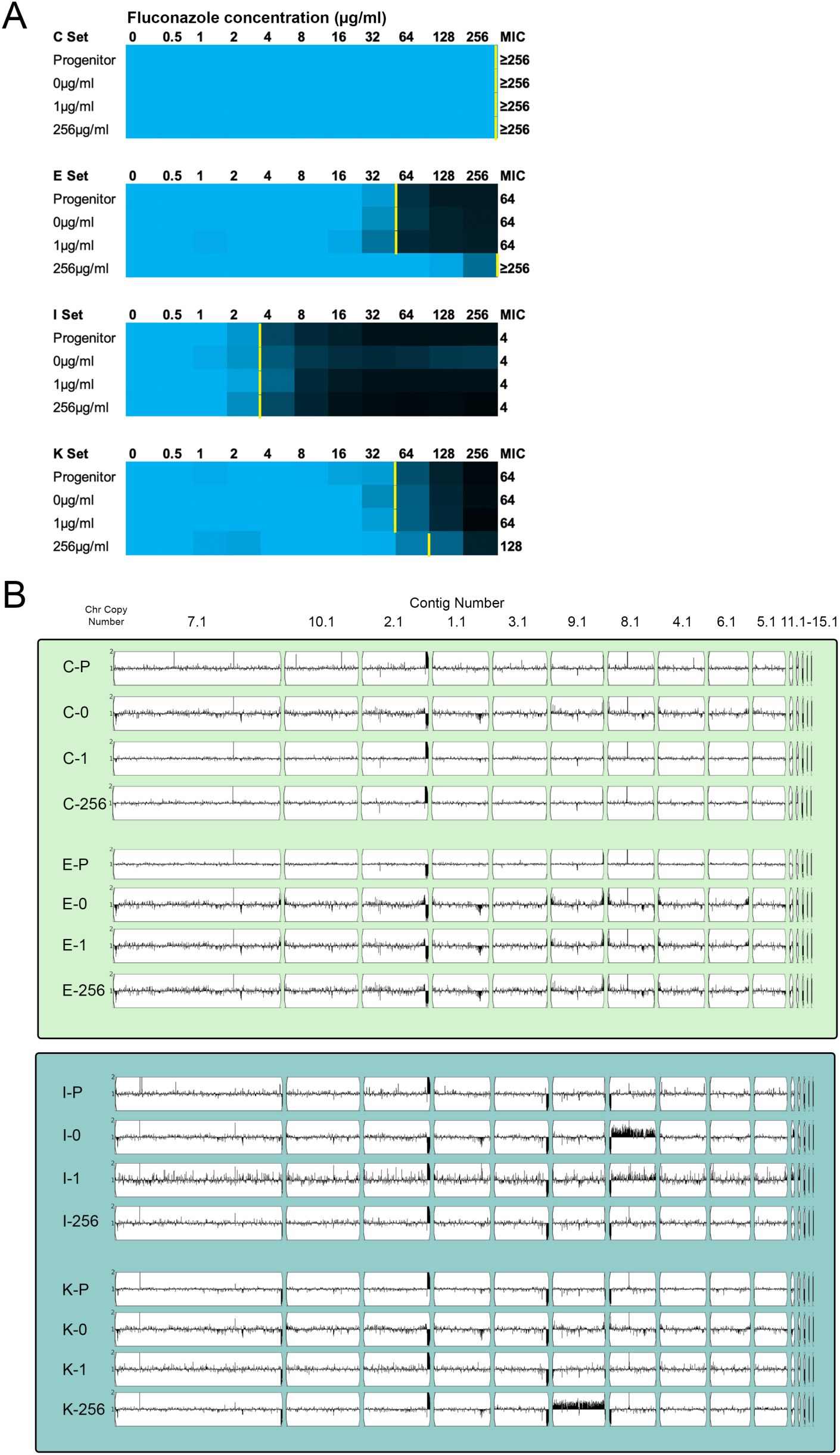
Determination of fluconazole MICs and genome karyotypes of *in vitro* evolution populations. (A) MICs were measured for all evolution lineages and representative evolution populations C, E, I and K from each evolution condition (0µg/ml, 1µg/ml, or 256µg/ml fluconazole (FLC)) are shown here below the progenitor data. MICs were measured by broth dilution assay in 0.5X dextrose YPAD with increasing concentrations of FLC. The MIC_50_ was defined as the lowest concentration of FLC that decreased the OD_600_ at 24 hr to less than 0.5 of the growth control without drug. Normalized growth is shown as a color scale where relative growth ≥0.8 is shown as blue and no growth is shown as black with scaled colors in between. The MIC_50_ for each is marked with a yellow bar. Normalized growth values shown are the mean of three replicates. (B) Whole-genome sequence data plotted using YMAP. Read depth was converted to copy number (Y-axis) and was plotted as a function of chromosome position using the B8441 Clade I reference genome for each progenitor and evolution population.

### *Candida auris* rapidly acquires aneuploidies and SNVs during *in vitro* evolution

Growth in fluconazole has been shown to promote whole genome ploidy increases in *C. albicans* [45]. To determine if *C. auris* changed ploidy during the short *in vitro* evolution experiment, we measured ploidy using flow cytometry. All the progenitors and evolved populations were haploid (Table S1). We also examined karyotypes via CHEF. Overall, the karyotypes of the evolved isolates were similar to the progenitors, but small changes in migration consistent with changes in repeat copy number and/or sub-telomeric loss were observed (Figure S3). To further examine CNVs, we performed Illumina WGS and mapped read depth using YMAP. Two evolved populations, I-0 and K-256, had detectable whole chromosome aneuploidies of chromosome 6 (contigs 8.1 and 12.1) and chromosome 5 (contig 9.1) (Figure 3B). CNVs were verified by read depth with IGV, and the average read depth across these chromosomes (∼1.3x) indicated that the aneuploid chromosome is only present in a subset of cells within the evolved populations (Figure 3B). Chromosome 5, which was amplified in K-256, contains several genes previously associated with drug resistance including *TAC1B.* The mechanism of selection for chromosome 6 aneuploidy (I-0) during *in vitro* evolution in the absence of drug is not immediately apparent, but it does have genes associated with drug resistance such as *B9J08_004113,* the homolog of *C. albicans MDR1,* suggesting that the aneuploidy, once selected, could influence antifungal resistance as well. Overall, this analysis shows that *C. auris* aneuploidies can arise rapidly (3 passages) and reach detectable levels in the population during *in vitro* evolution, both in the presence and absence of fluconazole. This suggests that aneuploidy may be a common mechanism of generating genome diversity in *C. auris*.

We next analyzed the WGS data for *de novo* SNVs by comparing the evolved populations with their respective progenitor isolate. We identified 98 SNVs causing either a moderate or high impact mutation that arose during the *in vitro* evolution experiments (Table S5). Of these 98 SNVs, we re-sequenced a subset of 28 SNVs spanning a range of allele frequencies with Sanger sequencing. Twenty-one of these SNVs were clearly visible in the Sanger sequencing traces. All seven remaining SNVs were at allele frequencies ≤0.2 suggesting that these alleles are below the level of detection of Sanger sequencing, but detectable in multiple reads with Illumina WGS (Table S5). The 98 SNVs arose in 83 different ORFs. To determine if gene functions were enriched in the SNVs that arose during *in vitro* evolution, we conducted gene ontology (GO) analysis using the *Candida* Genome Database. No GO terms were significantly (p<0.05) enriched. However, the strength of the GO term analysis was limited by the GO enrichment algorithm’s removal of multiple hits to the same ORF. We observed recurrent mutations in fifteen ORFs. Recurrent *de novo* mutations occurred both as the same SNV in multiple evolution populations (ex. S386P in B9J08_003450 in L-0 and L-1) and as different SNVs in the same ORF (ex. V742A and L760S in Tac1b in E-256 and K-256). Several of the recurrent mutations occurred in homologs of *C. albicans* genes related to nutrient acquisition, transcriptional regulation and regulation of the cell cycle as discussed below (Table S5).

### Rapid acquisition of recurrent mutations during *in vitro* evolution without changes in fluconazole resistance

Several recurrent mutations arose in independent lineages without accompanying changes in fluconazole resistance indicating that their selection during the *in vitro* evolution condition correlates with *in vitro* fitness more generally. In some cases, the *in vitro* evolution rapidly selected for alleles likely present at low levels within the progenitor obtained directly from the clinic. For example, both the N-0 and N-1 evolved populations had V406A missense mutations within ORF *B9J08_001332*, the ortholog of the transcription factor *PHO4*. The mutation was not detectable in the progenitor isolate, but the frequency of the mutation in both evolved populations was 1.0 indicating a very rapid sweep during the evolution experiment (Table S5). Similarly, an SNV in *B9J08_003619*, which encodes the ortholog of the *MED15* component of the mediator complex involved in transcription, increased from a frequency of 0.04 in the K-P clinical isolate to 0.79 in the K-0 evolved population and 0.29 in the K-1 evolved population (Table S5).

In other cases, multiple independent mutations arose within the same ORF in independent lineages. Most strikingly, nonsense mutations were observed in *IRA2,* the GTPase activator that negatively regulates *RAS1,* in three independent evolution populations, B-1, A-256A and C-256 (Table S1). The *IRA2* allele frequencies within the evolution populations were 0.2, 0.83, and 0.45 respectively. While these mutations map to two different ORFs (*B9J08_003924* and *B9J08_003925*) both annotated as *IRA2* homologs in the B8441 *C. auris* reference genome, we have several lines of evidence to support that these ORFs encode a single *IRA2* gene product (Figure S4A). First, the B8441 reference genome sequence contains an erroneous single base insertion that causes a frameshift that separates the two ORFs. This insertion was not observed with Sanger sequencing of B8441 and was absent from all other clade I sequences, indicating that it is a reference sequence error (Figure S4B). Furthermore, RNAseq data [46] was consistent with a single transcript. Based on the corrected ORF sequence, B-1 has a E677* nonsense mutation and A-256A and C-256 both acquired the same C1029* nonsense mutation. Additionally, while analyzing the *IRA2* mutations, we also determined that the B-P clinical isolate also had a third nonsense mutation, E1060* that was enriched during evolution experiment. The presence of this nonsense mutation in the original clinical isolate highlighted this gene as potentially important in regulating growth of *C. auris in vivo* as well.

The nonsense mutations in *IRA2* result in the deletion of the GTPase activating domain, so we hypothesized that these *IRA2* mutations would result in increased Ras signaling and potentially in increased growth *in vitro* [47, 48]. Therefore, we conducted growth curve analysis with three pairs of single colonies isolated from the evolved populations comparing the growth of those that contained the *IRA2* nonsense mutations with those that did not. No detectable differences in growth phenotypes were observed in the presence or absence of drug (Figure S4C). In addition to altered growth phenotypes in other species [47, 48], *IRA2* has been shown to be required for biofilm formation in *C. albicans* [49]. We hypothesized that the selective advantage of removal of *IRA2* function in our *in vitro* evolution experiment may be due to decreased biofilm formation resulting in more planktonic cells available for transfer to the next passage. Therefore, we measured biofilm formation in the progenitor isolates and representative single colonies with and without *IRA2* stop mutations using a crystal violet assay. We used *C. albicans* SC5314 as a known biofilm-forming control. As expected, *C. albicans* formed a robust biofilm in YPAD, but it did not grow well or form a biofilm in YPAD+128µg/ml fluconazole (Figure S4D). *C. auris* isolates formed biofilms in both conditions, although the biofilms were weaker than *C. albicans*. Although there was isolate-to-isolate variability in biofilm formation, we did not observe any significant differences between the progenitors and evolved isolates with or without *IRA2* nonsense mutations (Figure S4D). Overall, our results demonstrate that nonsense mutations in *IRA2* are repeatedly selected for during *in vitro* evolution in the presence of fluconazole and truncation of *IRA2* can also be observed directly in clinical isolates. Even though we did not observe significant differences in growth rates of individual colonies, it is possible that in direct competition and/or in environments with nutritional limitations or other constraints, inactivation of *IRA2* has a selective advantage both *in vitro* and *in vivo*.

### Multiple mechanisms to acquire fluconazole resistance occur simultaneously in *C. auris* populations

We only observed increases in MIC when cells were grown in 256µg/ml fluconazole (Figure 3A and Table S1), so we first determined whether this high concentration of fluconazole impacted the number and/or the gene targets of the moderate and high impact SNV mutations. In the combined set of seventeen evolution populations from each condition, the number of *de novo* SNVs acquired in 0µg/ml, 1µg/ml and 256µg/ml fluconazole evolution conditions was similar (31, 30 and 37 SNVs, respectively). GO analysis identified significant (p<0.05) enrichment of gene functions involved in protein tyrosine kinase activity and ion binding in the 256µg/ml fluconazole condition. Within the group of ion binding ORFs, the number of zinc binding transcription factors was striking, including multiple independent SNVs in *TAC1B* and the *C. auris* homolog of *UPC2* in the two populations that acquired a significant increase in fluconazole MIC during the evolution experiment (E-256 and K-256) (Figure 3 and Table S5). Numerous mutations in *TAC1B,* the transcription activator of Cdr efflux pumps, have previously been associated with fluconazole resistance in *C. auris* [11]. In other *Candida* species, *UPC2* homologs are key regulators of antifungal drug resistance and gain-of-function mutations in *UPC2* increase transcription of ergosterol biosynthesis genes [50–52]. These data highlight the speed (∼30 generations) in which *de novo UPC2* and *TAC1B* alleles can arise and expand within a population.

The *TAC1B*^V742A^ mutation arose in the E-256 lineage from clade I and reached an allele frequency of 0.79 (Table S5). Interestingly, the *TAC1B*^V742A^ variant along with an increase in fluconazole MIC from 64 to >256µg/ml in the E-256 population (Figure 3). No other moderate or high impact mutations or aneuploidies were observed in the E-256 population (Table S5 and Figure 3), indicating that the *TAC1B* missense variant is likely driving the 4-fold increase in MIC. Unlike the E-256 population, the mutational landscape in the K-256 population which increased in fluconazole MIC from 64 to 128µg/ml fluconazole was much more complicated. The K-256 population acquired the *TAC1B*^L760S^ mutation at a frequency of 0.26, and two different missense mutations in *UPC2*, C444Y and A506V, arose at allele frequencies of 0.14 and 0.15 respectively during the *in vitro* evolution experiment (Table S5). In addition to the *TAC1B* and *UPC2* SNVs, the K-256 population had 20 other detectable moderate and high impact SNVs and an aneuploidy on contig 9.1 suggesting that multiple mechanisms of resistance are arising simultaneously in subsets of the K-256 population.

### Mutational Landscape within Evolution Populations and Evidence of Clonal Interference

To further explore the mutational landscape of the evolution lineages, we isolated single colonies from selected populations and conducted Illumina WGS. We selected populations for single colony analysis based on SNVs (e.g. two different *IRA2* stop alleles within the B-1 population) and presence of aneuploidies within the population (I-1 and K-256 populations). We characterized and sequenced six individual colonies from the B-1, I-1, K-1 and O-1 evolution populations and twenty-four single colonies from the K-256 population (Table S1). These single colony experiments highlighted extensive phenotypic and genotypic variability within the evolution populations, especially in the K-256 lineage (Figure 4 and Figure S5).

**Figure 4:**
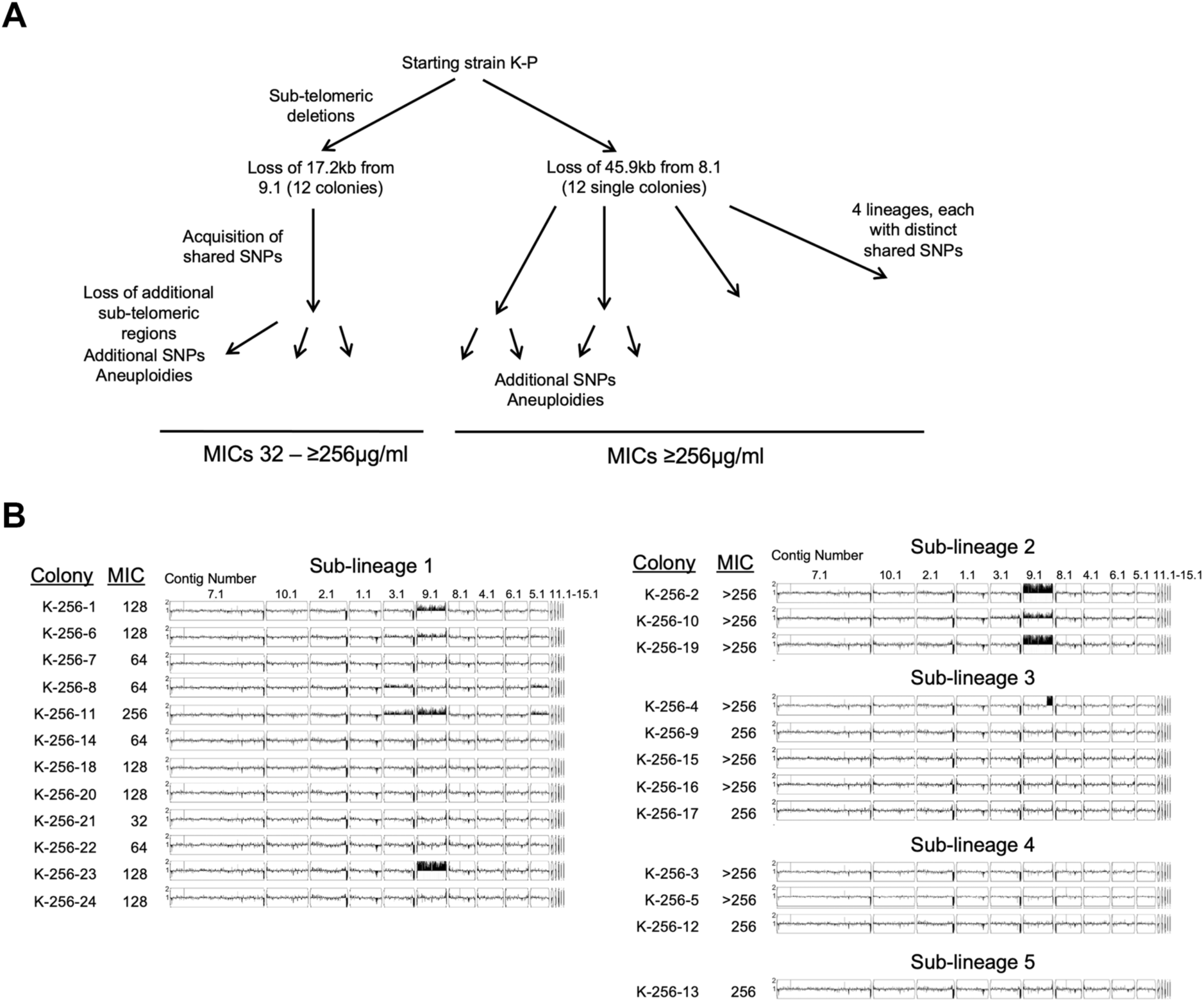
Genomic and phenotypic diversity within the K-256 evolution population. (A) Schematic of shared sub-telomeric deletions and SNVs to identify the five sub-lineages of the K-256 evolution population. (B) MICs were measured for each single colony from the K-256 population. MICs were measured by broth dilution assay in 0.5X dextrose YPAD with increasing concentrations of FLC. The MIC_50_ was defined as the lowest concentration of FLC that decreased the OD_600_ at 24 hr to less than 0.5 of the growth control without drug. Whole-genome sequence data plotted using YMAP. Read depth was converted to copy number (Y-axis) and was plotted as a function of chromosome position using the B8441 Clade I reference genome for each single colony.

One striking result was the detection of additional sub-telomeric deletions within single colonies of the evolution populations (Table S1 and Table S2). One colony from each set of six from the B-1 and I-1 populations underwent an additional sub-telomeric deletions. In the K lineages, the sub-telomeric regions showed even greater rates of change. Three K-1 colonies underwent 1-2 additional sub-telomeric deletions. Within the K-256 single colonies, all 24 single colonies underwent additional sub-telomeric deletions of 17.2kb to 45.9kb that were not apparent in the original population (Figure 4A and Table S2). Furthermore, among the colonies that lost 17.2kb (from contig 9.1), two colonies, K-256-1 and K-256-11, each had an additional deletion of 1.1kb and 2.6kb, respectively (Table S2). Overall, these results show highly dynamic sub-telomeric regions in the presence of fluconazole where changes can become apparent on the timescale of three passages totaling approximately 30 generations.

Together with chromosome 5 aneuploidy in a subset of the population and multiple SNVs in fluconazole resistance genes, we hypothesized that the K population had simultaneously evolved into multiple sub-lineages with clonal interference preventing a sweep of the population due to competition between mutants with enhanced fluconazole resistance. To test this hypothesis, we tracked the different evolutionary trajectories within the K-256 population using the unique sub-telomeric deletions and SNVs found in each single colony. We were able to divide the K-256 population into five distinct sub-lineages and analyzed the SNVs and aneuploidies in these isolates to identify genetic determinants of resistance in the sub-lineages. The K-256 population had a fluconazole MIC of 128µg/ml, while the single colonies had fluconazole MICs ranging from 32µg/ml up to >256µg/ml indicating a high level of heterogeneity within the population (Figure 4B), while the MICs of single colonies from the I-1 and K-1 populations were similar to each other and to the population (Table S1).

Sub-lineage 1 had the greatest diversity in MIC values and several aneuploidies. We observed several SNVs shared among sub-lineage 1 isolates (Table S5), but none of the SNVs identified have been previously linked to azole resistance. Instead, aneuploidy seems to be the driving force for fluconazole resistance in sub-lineage 1. Colonies K-256-1, K-256-6, K-256-11 and K-256-23 had increased copy number of contig 9.1/Chromosome 5 and corresponding increased fluconazole MIC values (Figure 4B). This chromosome contains several genes potentially associated with azole resistance including *TAC1A* and *TAC1B* as well as *NPC1, ERG9* and *ERG13*. Colony K-256-11 had the highest MIC_50_ (256µg/ml fluconazole) in sub-lineage 1 and had increased copy number of two additional contigs (3.1 and 5.1) which both map to Chromosome 3. Interestingly, Chromosome 3 contains the *ERG11* gene which is the target of fluconazole.

For sub-lineages 2 through 5, all colonies had fluconazole MICs ≥256µg/ml, which was the limit of detection. Sub-lineage 2 had increased copy number of contig 9.1/Chromosome 5 as well as the *UPC2*^A506V^ variant (Figure 4B and Table S5). Sub-lineage 3 shared the *UPC2*^C444Y^ variant among all colonies suggesting a possible mechanism for increased resistance (Table S5). Intriguingly, one colony from sub-lineage 3, K-256-4, also contained a novel segmental aneuploidy on contig 9.1 that includes the *TAC1B* gene (Figure 4B). Sub-lineages 4 and 5 had high MICs, but neither had detectable aneuploidies or SNVs previously associated with resistance, opening up the possibility of new resistance mechanisms. For example, Sub-lineage 5 had missense mutations in the *C. auris* homologs of *DCR1*, the Dicer ribonuclease III enzyme, and *TRA1*, a histone acetyltransferase suggesting that alterations in the regulation of gene expression are increasing fluconazole resistance in colony K-256-13 (Table S5). Together, these results indicate that multiple paths to increased resistance including missense SNVs and whole chromosome and segmental aneuploidies can occur very rapidly in just three passages within a single evolution population.

### Clinical Isolate K Has Elevated Mutation Rates

In examining the complete set of mutation data, we were struck by the mutational diversity that arose from the K-P progenitor relative to other isolates (Figure 5A). Indeed, a majority of all SNVs were from the K-P progenitor: 20/31SNVs identified in the 0µg/ml fluconazole evolution populations were from the K-0 population and only 11 were from the other 16 progenitors. Similarly, 20/30 SNVs and 23/37 SNVs from the 1µg/ml and 256µg/ml fluconazole evolution populations were from K-1 and K-256. Given the high rate of acquisition of SNVs in the *in vitro* evolution experiment relative to the other progenitor isolates, we decided to measure the rate of loss of *URA3* function as a marker of mutational rate using 5-fluororatic acid (5-FOA) selection. We compared loss rates of *URA3* per cell division using a fluctuation assay with K-P as well as two other closely related clade IV isolates H-P and I-P. The progenitor isolate K-P had approximately a 10-fold higher 5-FOA resistance rate with loss of function mutations arising at 2.1×10^-7^ mutations/cell division while H-P and I-P had loss rates of 4.4×10^-8^ and 1.3×10^-8^ mutations/cell division, respectively (Figure 5B).

**Figure 5:**
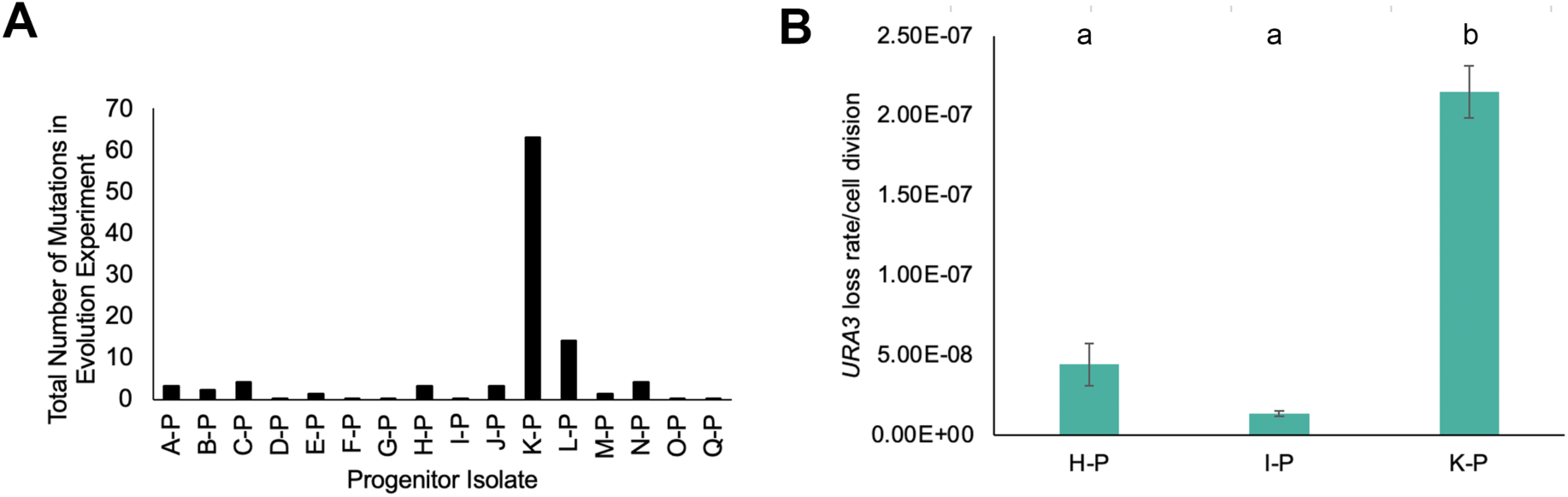
Elevated mutation rates in K lineages. (A) Total number of SNVs over the course of the evolution experiment for all three conditions for each progenitor. Moderate and high impact mutations were identified by Mutect2 by comparing each ending evolution population to the progenitor strain. (B) Progenitor isolates were grown overnight in YPAD media, then diluted with water and plated onto YPAD for total cell counts and 5-FOA plates to select for *URA3* loss. Colony counts were used to calculate the rate of loss per cell division. Data are mean ± SEM calculated from at least three biological replicates, each with eight cultures per conditions. Significant differences are indicated with letters (Ordinary one-way ANOVA, p<0.0001; Tukey multiple comparison, p<0.001 for letter difference).

To determine possible genetic determinants of the high mutation rate of K-P relative to H-P and I-P, we identified SNVs unique to K-P. There were ten missense or nonsense impact variants found in K-P compared to isolates H-P and I-P (Table S4). As expected, one of these variants was the *ERG11^Y132F^* mutation that likely confers the higher starting fluconazole resistance of K-P relative to H-P and I-P. Strikingly, one of the variants was a A216V missense mutation in *B9J08_004425* which encodes the *C. auris* homolog of the mismatch DNA repair component *MLH1.* These data suggest a possible alteration in mismatch repair is associated with the higher mutational rate in original clinical isolate (K-P). To the best of our knowledge, this is the first description of a mutator phenotype in a *C. auris* clinical isolate.

## Discussion

### High rates and stability of acquired antifungal drug resistance

Through *in vitro* evolution of a set of seventeen new clinical isolates of *C. auris* from clades I and IV we determined that *C. auris* can rapidly undergo genomic and phenotypic changes including aneuploidy, loss of sub-telomeric regions, acquisition of *de novo* point mutations, and increases in antifungal MIC values. In just 30 generations over 9 days, two different progenitor isolates (E-P and K-P) acquired two-fold or greater increases in fluconazole MIC_50_ values when grown in 256µg/ml fluconazole (Figure 3). Previously published *in vitro* evolution experiments with *C. auris* have suggested that fluconazole resistance can develop rapidly. Bing *et al.* observed increased fluconazole MICs up to 128µg/ml after 18 days in a clade I strain [13], and Carolus *et al.* detected increased fluconazole MICs up to 32µg/ml as early as day 13 in a clade II strain [16]. Our experiments demonstrate rapid evolvability of clinical isolates from both clades I and IV over even shorter timescales. Importantly, acquisition of resistance was much faster than loss of resistance. We observed little to no fitness cost associate with resistance as all clinical isolates with high starting MICs maintained resistance in the absence of drug. Similarly, even though we did not include echinocandin treatment, the *FKS1* mutations associated with echinocandin resistance were maintained at an allele frequency of 1.0 throughout the evolution experiment. This means that strains can accumulate resistance to different classes of antifungal agents. Once resistance is acquired, it is maintained in a *C. auris* population even in the absence of selective pressure for an extended period. If the drug treatment is stopped or switched, antifungal resistance is unlikely to be lost which sets a pathway towards the evolution of multidrug-resistant and pan-resistant isolates of *C. auris* [9, 53].

### Multiple mechanisms of resistance and tolerance

Our mutational spectrum results suggest that the probability of evolution of multidrug-resistant and pan-resistant isolates is also elevated in *C. auris* because of the large number of different mechanisms that promote acquired drug resistance. Strikingly, many of these different mechanisms promoting fluconazole resistance arose simultaneously in a single population (K-256) further emphasizing a high likelihood of clonal interference where multiple beneficial mutations are selected for simultaneously [48]. This mutator isolate increased its fluconazole MIC_50_ to >256µg/ml, and resistance occurred in at least five different evolutionary pathways concurrently (Figure 4 and Table S5). Individual isolates from sub-lineage 1 had an increased copy number of Chromosome 5 and Chromosome 3. Sub-lineages 2 and 3 had distinct *UPC2* mutations as well as increased copy number of all or part of Chromosome 5. Amplification of *TAC1B* and *ERG11* has been observed in previous evolution experiments [13, 16] suggesting it is a recurring mechanism in antifungal drug resistance acquisition. Perhaps most intriguingly, *s*ub-lineages 4 and 5 had high MICs, but none of these single colonies had detectable aneuploidies or SNVs previously associated with resistance indicating that there are additional novel mechanisms of fluconazole resistance. For example, alteration of gene expression may occur in mutants that acquired missense mutations in the *C. auris* homologs of *DCR1*, the Dicer ribonuclease III enzyme, and *TRA1*, a histone acetyltransferase (Table S5).

Zn2-Cys6 zinc finger transcription factors were enriched in the evolution mutation set with recurrent mutations in *TAC1B* as well as the *C. auris* homologs of *C. albicans UPC2, ZCF18,* and *ZCF22* genes suggesting that these transcription factors regulate cellular processes related to fluconazole import, export or targeting effects. We uncovered new *TAC1B* and *UPC2* alleles not previously associated with increased fluconazole resistance (Table S5). Among clinical isolates, mutations in *TAC1B* are common, and five of the six clade I isolates included in this collection had the *TAC1B*^A640V^ allele seen in other clade I clinical isolates with high levels of resistance [11]. The *TAC1B*^V742A^ and *TAC1B*^L760S^ alleles that arose during the evolution experiments have not been previously identified, suggesting that the range of possible alleles that can be mutated in *TAC1B* to result in activation is even higher than previously appreciated. In addition to *TAC1,* several zinc cluster transcription factors, including *UPC2,* have been identified as essential for fluconazole resistance in *C. albicans,* and activating mutations in *UPC2* elevate fluconazole resistance by increasing transcription of the ergosterol biosynthesis pathway [54]. A *UPC2* missense mutation in a single clade I isolate had previously been shown to correlate with a 2-fold increase in MIC_50_ [11]. While the mechanistic role of *UPC2* has not been characterized yet in *C. auris,* the identification of *UPC2* mutations associated with resistance in two independent evolution experiments and in two different sub-lineages of K-256 indicates that *UPC2* is important in regulating resistance to fluconazole in *C. auris*.

In addition to fluconazole resistance (both pre-existing and evolved), we also characterized the first examples of fluconazole tolerance in *C. auris*. Tolerance did not increase during evolution, but several clinical isolates had high levels of initial tolerance. Our comparative genomics identified possible genetic factors contributing to high tolerance, including *ERG3*. Genetic background has previously been shown to be a determinant of tolerance in *C. albicans* [23], but there is a limited understanding of the specific genes and alleles associated with tolerance in any Candida species. *ERG3* is involved in the production of toxic by-products resulting from blockage of *ERG11* by azoles such as fluconazole such that changes in *ERG3* function may allow slow growth in the presence of fluconazole [55]. *ERG3* mutations have also previously been associated with resistance to polyenes and echinocandins [16, 56] and have been shown to drive cross-resistance to multiple antifungals [22]. In work by Carolus *et al.,* an *ERG3* truncation mutation arose during *in vitro* evolution of *C. auris* in the presence of amphotericin B, and this mutation was also associated with elevated fluconazole resistance [16]. Combination therapies show promise for treating pan-resistant isolates of *C. auris* [53]. However, when using antifungal resistance genes as biomarkers for resistance clinically, it will be important to consider the multifactorial effects of these mutations, especially as combination therapies are considered.

Most of the mutations that occurred during the *in vitro* evolution experiment in fluconazole were SNVs while a minority of isolates acquired aneuploidies. This is in contrast to rapid evolution of fluconazole resistance in *C. albicans* where CNVs in the form of aneuploidy and segmental amplifications are the dominant type of resistance mechanism observed after short-term *in vitro* evolution [19]. One possible explanation for the differences observed is due to the diploid genome in *C. albicans vs* the haploid genome in *C. auris*. SNVs in a diploid genome must be dominant to have a phenotypic effect thus masking the effects of recessive mutations. In a haploid genome, only one copy of the gene is present such that SNVs can have immediate phenotypic impact [20, 21].

### Genome instability in sub-telomeres

We observed rapid and dynamic genome instability of the *C. auris* sub-telomeres *in vitro* and *in vivo*. The haploid genome structure in *C. auris* also increases the likelihood that there will be phenotypic consequences associated with loss of sub-telomere regions. Every clinical isolate in our collection had at least one sub-telomeric deletion compared to the reference genome B8441 (Table S2). Strains within a clade are similar at the sequence level with few SNVs (Figure 1), but they differ with regards to the gene content at the sub-telomeres. Prior work has shown that relative to the reference genome (B8441/clade I), clade II strains lost adhesins and cell wall proteins from sub-telomeric regions [5]. However, our results demonstrate that this is a widespread phenomenon that results in genotypic diversity both between clades and within a given clade of *C. auris* and that adhesins and cell wall proteins are a minority of the ORFs that are lost in the sub-telomeres. Instead, GO term analysis of the annotated gene functions of ORFs within the deleted sub-telomeric regions showed significant enrichment for transmembrane transporter activity indicating that import and export of nutrients such as iron and glucose are likely affected as well as import and export of antifungals. When comparing H-P (high fluconazole tolerance) and K-P (fluconazole resistance and high mutability) to other clade IV strains, including I-P from the same patient, they had lost an additional three ORFs on Contig 7.1. One of these ORFs is annotated as a *SIT1* homolog in *C. albicans* with roles in iron transport and another is annotated as the homolog of the plasma membrane transporter *DAL9* in *C. albicans* which is similar to *DAL5* in *S. cerevisiae*. *ScDAL5* null mutants have more than twice the normal deposition of chitin [57]. Given the role of excess chitin production in enhancing antifungal drug resistance in *C. albicans* [58], it is tempting to speculate that the loss of this sub-telomeric gene may also be a contributing factor to fluconazole tolerance/resistance and/or echinocandin resistance.

In addition to pre-existing sub-telomeric differences between clinical isolates, we also observed loss of sub-telomere regions during *in vitro* evolution. Closer examination of these loss events *in vitro* showed the loss of sequence in one of the sub-telomere regions, but no additional copy number of any other sub-telomeres suggesting that loss occurs due to a recombination method that does not involve repair via another chromosome such as break-induced replication (BIR) [19]. Sub-telomere regions are dynamic in other fungal species as well. In *C. albicans*, the sub-telomere regions contain a set of *TLO* genes that are expanded compared to other closely related *Candida* species and have elevated mutation and LOH rates [59]. Non-reciprocal recombination events including crossovers and gene conversions also occurred at the *C. albicans* telomeres during *in vitro* evolution experiments resulting in changes in *TLO* genes [60]. However, in contrast to what we observed in *C. auris*, changes to the *TLO* genes in *C. albicans* resulted in acquisition of a different *TLO* or formation of a hybrid gene rather than reduction in *TLO* number [60].

### Genetic background/mutator phenotypes

One of the most striking results was the importance of genetic background on the spectrum of acquired mutations. The frequency of SNVs varied significantly depending on the progenitor isolate with one progenitor isolate, K-P, accounting for more than half of all the SNVs identified in the entire experiment. K-P also had elevated rates of 5-FOA resistance as an indicator of mutation rate (Figure 5). Estimated rates of SNP acquisition in *C. auris* varies dramatically among published studies from 5.75 SNPs per genome per year [61] to more than 400 SNPs per year [62] with genetic variability observed both with hosts and within healthcare facilities suggesting that clinically important evolution can occur both in the environment and *in vivo* [62, 63]. Differences in estimated mutation rate could be explained by different stringency in bioinformatic approaches, differences in estimates of the effective population size, growth rates, or selection pressure due to antifungal drug treatment [61]. However, differences in rates of mutation during outbreaks also may be explained by biologically important differences in the genetic background such as alterations to DNA repair proteins as seen in the K-P isolate. The K-P isolate described here is the first known example of a mutator phenotype in *C. auris* (Figure 5). Elevated mutation rates have been observed in other fungal pathogens including *Cryptococcus* species mutator alleles in DNA repair pathway genes and activation of transposase activity enhance within-host adaptation and evolution of antifungal resistance [64–67].

### Tracing the evolutionary pathway of multi-drug resistance

The first isolate of *C. auris* identified in 2009 was drug-sensitive as is the reference genome strain B8441 first isolated in Pakistan in 2010 [4], but since then rapid acquisition of multidrug resistance has occurred worldwide. Multidrug resistant isolates can spread from one patient to another [9], and molecular epidemiological tracking has shown that resistance can arise rapidly. For example, genomic evaluation of nosocomial *C. auris* transmission in the Chicago metro region indicated that all isolates likely trace back to a single introduction of a drug-sensitive clade IV isolate followed by patient-to-patient transmission and genomic diversification [68]. Within the region, fluconazole resistance has evolved independently multiple times [68]. By combining the genomic analysis and evolutionary insights from this study with prior molecular epidemiological tracking of *C. auris*, we put together a model of how multidrug resistance and a high mutation rate evolved in an individual patient over the course of infection (Figure 6). For this model, we focused on the three clade IV clinical isolates from a single patient in Illinois where echinocandin resistance, fluconazole tolerance and fluconazole resistance evolved during the course of infection (Figure 4, Table S1 and [33]). By tracking sub-telomeric deletions and comparative genomic analysis of SNVs (Table S1 and Table S4), we reconstructed the possible evolutionary path of the clinical isolates and highlight potential evolutionary tracts that are clinically relevant for the evolution of multi-drug resistance *in vivo* and further increases in MICs during *in vitro* evolution (Figure 6). An *FKS1^S639P^* mutation in the progenitor of H-P and K-P was maintained even as large sub-telomeric deletions and other SNVs accumulated both *in vivo* and *in vitro* (Figure 6A). In this echinocandin-resistant background, both fluconazole tolerance and moderate fluconazole resistance evolved independently within the same patient following treatment with posaconazole and fluconazole [33]. H-P acquired tolerance and a set of mutations including an *ERG3* missense mutation. While in the K-P lineage, acquired mutations included missense alleles in *ERG11* and *MLH1* correlated with moderately elevated fluconazole resistance and a higher mutation rate. Together, these mutations allowed further acquisition of SNVs and aneuploidies during only three passages of *in vitro* evolution that elevated fluconazole resistance through many different mechanisms to strengthen the multi-drug resistance of this isolate (Figure 6B). Overall, the genome analysis of clinical isolates and *in vitro* evolution experiments here provide insights into the high stability and high rates of acquisition of antifungal resistance of *C. auris* that allow evolution of pan-resistant, transmissible isolates in the clinic.

**Figure 6:**
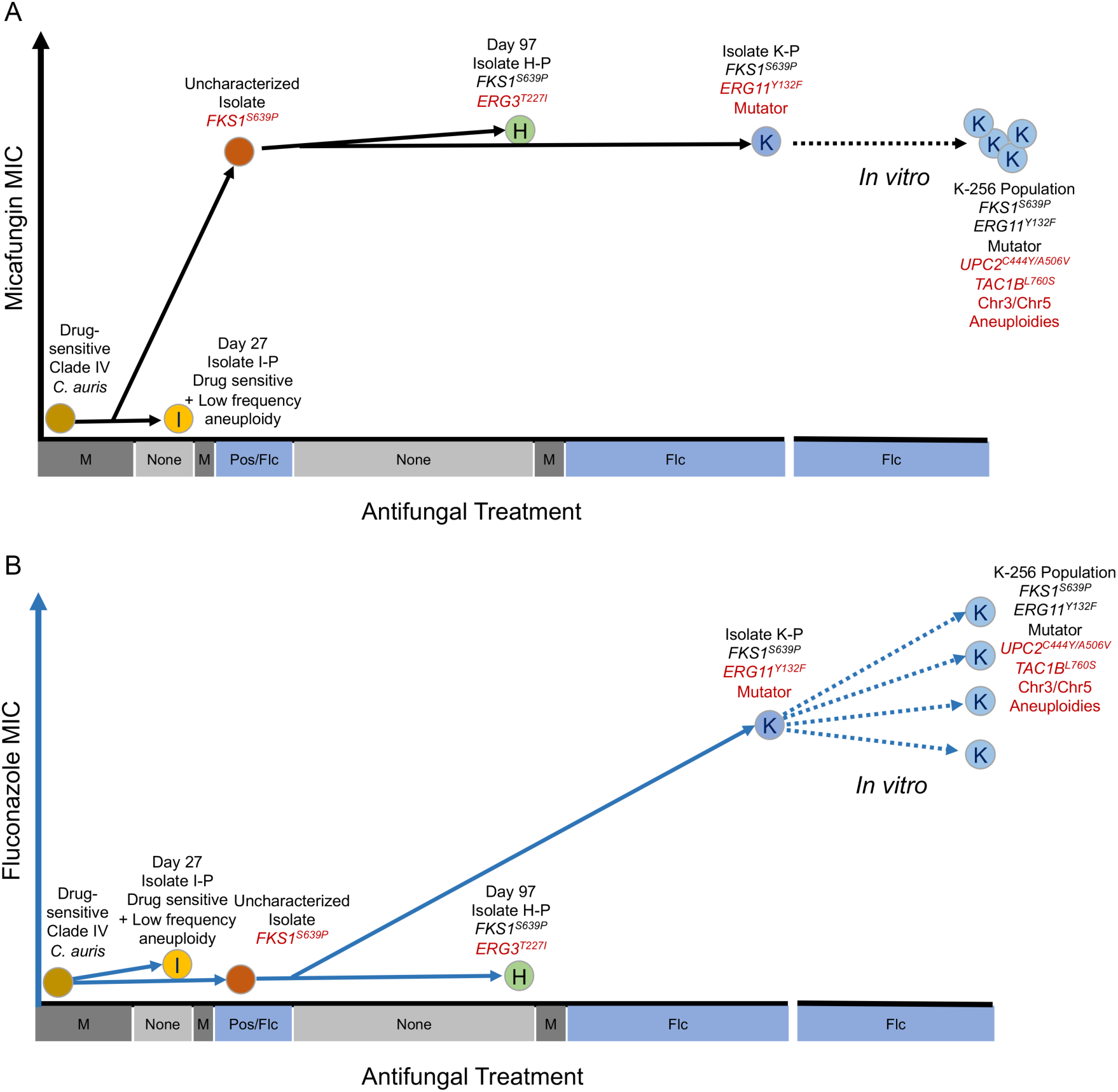
Schematic of evolutionary path of rapid acquisition of multidrug resistance. (A) Evolution of micafungin resistance associated with acquisition of *FKS1^S639P^* occurred during the course of infection and treatment with micafungin and two different azoles (posaconazole and fluconazole) in a single patient. The *FKS1^S639P^* allele was maintained both *in vivo* and during *in vitro* evolution in the absence of echinocandins. (B) Evolution of fluconazole tolerance (H) associated with *ERG3^T227I^* and moderate fluconazole resistance (K) associated with *ERG11^Y132F^* occurred in parallel from a shared echinocandin resistant progenitor. K also acquired a mutator phenotype and rapidly evolved to higher resistance to fluconazole through a variety of mechanisms during *in vitro* evolution.

## Materials and Methods

### Yeast isolates and culture conditions

All isolates used in this study are described in Table S1. Clinical *C. auris* isolates were obtained from the Fungus Testing Laboratory at the University of Texas Health Science Center at San Antonio. All isolates were identified as *Candida auris* by DNA sequence analysis of the ITS region and large ribosomal subunit gene and phenotypic characteristics as previously described [33]. Isolates were stored at -80° C in 20% glycerol. Isolates were cultured at 30°C in YPAD medium (10g/L yeast extract, 20g/L bactopeptone, 0.04g/L adenine, 0.08g/L uridine, 20g/L dextrose with 15g/L agar for plates) unless otherwise specified. For selection for loss of *URA3* function, cells were grown on SDC+ 1g/L 5-FOA supplemented with 0.1g/L uridine and 0.1g/L uracil.

#### Antifungals and Clinical Susceptibility Testing

Susceptibility testing in Table S3 was performed at the Fungus Testing Laboratory at the University of Texas Health Science Center according to CLSI M27 broth microdilution methods [69]. Antifungal powders were purchased from SigmaAldrich (St. Louis, MO, USA; amphotericin B, caspofungin, micafungin, fluconazole, posaconazole, and voriconazole) or were provided by Astellas Pharmaceuticals (Northbrook, IL, USA). The concentration ranges tested were 0.015 to 8µg/ml for caspofungin and micafungin, 0.03 to 16µg/ml for amphotericin B, isavuconazole, posaconazole, and voriconazole, and 0.125 to 64µg/ml for fluconazole. The MIC_50_ for the echinocandins and azoles was the lowest concentration that resulted in at least 50% growth inhibition compared to the growth control after 24 hours of incubation at 35°C. For amphotericin B, the MIC was the lowest concentration that resulted in 100% inhibition of growth. The CLSI quality control isolates *Candida parapsilosis* ATCC 22019 and *C. krusei* ATCC 6258 were also included.

### *In vitro* evolution experiment

Seventeen *Candida auris* clinical isolates (A-P through Q-P) were plated on YPAD and incubated for 48 hours at 30°C. Cells were diluted into YPAD and the optical density (OD_600nm_) from a BioTek Epoch spectrophotometer was adjusted to 1.0. A 1:1000 dilution was made into three experimental groups: 1) YPAD, 2) YPAD with 1µg/ml fluconazole, and 3) YPAD with 256µg/ml fluconazole in deep-well 96-well plates. Plates were sealed with Breath EASIER tape (Electron Microscope Science) and placed in a humidified chamber for 72 hours at 30° C. Every 72 hours, cells were resuspended and transferred 1:1000 into fresh YPAD with the indicated concentration of fluconazole. After three transfers, cells were collected for DNA isolation, MIC and CHEF analysis, and for storage at -80°C. The evolution populations are named after their starting progenitor isolate and the drug concentration of the evolution experiment (ex. K-256). Two separate A-256 populations were evolved, so these populations are names A-256A and A-256B.

### Microdilution minimum inhibitory concentration (MIC)

The fluconazole MIC_50_ for all clinical and evolved isolates was measured using a microwell broth dilution assay in Table S1, Figure 3 and Figure 4. Isolates were inoculated from frozen stocks into YPAD and grown for 16-18 hours at 30°C. Cells were diluted in fresh 0.05X dextrose YPAD to a final OD_600nm_ of 0.01, and 20µl of this dilution was inoculated into a 96-well plate containing 180µl of a 0.5X dextrose YPAD with a 2-fold serial dilution of the antifungal drug or a no-drug control. Final concentrations of fluconazole tested ranged from 0.5µg/ml to 256µg/ml. Cells were incubated at 30°C in a humidified chamber and OD_600nm_ readings were taken at 24 and 48 hours post inoculation with a BioTek Epoch spectrophotometer. The MIC_50_ of each of the isolates was determined as the concentration of antifungal drug that decreased the OD_600nm_ by ≥50% of the no-drug control at 24 hours. The SMG (supra-MIC growth) was calculated as the average OD_600nm_ for all wells above the established MIC_50_ for that isolate/lineage at 24 hours normalized to the no-drug control [23, 24].

### Growth Curve Analysis

Isolates were inoculated from frozen stocks into YPAD and grown for 16hr at 30°C. Cells were diluted in fresh YPAD to a final OD_600nm_ of 0.01, and 10µl of this dilution was inoculated into a 96-well plate containing 190µl of YPAD with or without the indicated concentration of fluconazole. Cells were grown at 30°C in the BioTek Epoch with shaking (256 rpm) for 48hr with OD_600nm_ readings every 15 minutes. Growth curves were conducted in biological triplicate in three separate experiments. Calculation of summary statistics was conducted using the R package Growthcurver using standard parameters [70].

### Biofilm Assay

Isolates were patched from frozen stocks to YPAD plates and incubated for 72hr at room temperature. Cells were transferred to YPAD and diluted to an OD_600nm_ of approximately 0.05. 20µl of this dilution was added to 1ml YPAD with or without the indicated concentration of fluconazole. In triplicate, 200µl of this mixture was added to a polystyrene plate. Plates were covered with Breath EASIER tape (Electron Microscope Science) and placed in a humidified chamber for 72 hours at 30°C. To stain the biofilm, the planktonic cells were removed, then each well was washed 3X with sterile water prior to adding 200µl 0.1% crystal violet for 15 min at room temperature. The stain was removed, then each well was washed 3X with sterile water. Plates were allowed to dry for 15 min at room temperature, then 200µl solubilization solution (10% acetic acid, 20% methanol) was added to each well. Following incubation for 10 min at room temperature, OD_600nm_ was measured with a BioTek Epoch spectrophotometer.

### Contour-clamped homogenous electric field (CHEF) electrophoresis

Sample plugs were prepared as previously described [71]. Briefly, cells were suspended in 300µL 1% low-melt agarose (Bio-Rad) and digested with 1.2mg Zymolyase (US Biological) at 37°C for 16 hours. Plugs were washed twice in 50 mM EDTA and treated with 0.2 mg/ml proteinase K (Alpha Asar) at 50°C for 48 hours. Chromosomes were separated in a 1% Megabase agarose gel (BioRad) in 0.5X TBE using the CHEF DRIII Pulsed Field Electrophoresis System under the following conditions: 60s to 120s switch, 6 V/cm, 120° angle for 36hrs followed by 120s to 300s switch, 4.5 V/cm, 120° angle for 8 hours. CHEF gels were stained with ethidium bromide and imaged with the GelDock XR Imaging system (BioRad).

### Illumina whole genome sequencing

Genomic DNA was isolated using a phenol-chloroform extraction as described previously [14]. Libraries were prepared using either the Illumina Nextera XT DNA Library Preparation Kit or the Nextera DNA Flex Library Preparation Kit. Adaptor sequences and low-quality reads were trimmed using Trimmomatic (v0.33 LEADING:3 TRAILING:3 SLIDINGWINDOW:4:15 MINLEN:36 TOPHRED33) [72]. Trimmed reads were mapped to the *C. auris* reference genome (GCA_002759435.2_Cand_auris_B8441_V2) from the National Center for Biotechnology Information (https://www.ncbi.nlm.nih.gov/assembly/GCA_002759435.2/). Reads were mapped using BWA-MEM (v0.7.12) with default parameters {Li, 2013}. PCR duplicated reads were removed using Samtools (v0.1.19) [73], and realigned around predicted indels using the Genome Analysis Toolkit (RealignerTargetCreator and IndelRealigner, v3.4-46) [74].

### Phylogenomic analysis

Phylogenomic analysis was conducted for the starting clinical isolates (Table S1 and Figure 1). Variants were generated using realigned, sorted BAM files with the Genome Analysis Tool Kit (HaplotypeCaller, v3.7-0-gcfedb67) with the ploidy parameter of 1 [74]. Variants were combined using GATK’s CombineGVCFs and genotyped using GATK’s GenotypeGVCF to generate a single VCF file containing all identified variants. Variants were sorted as either SNVs or INDELs using GATK’s SelectVariants, and low-quality SNVs were removed using GATK’s VariantFiltration using the following parameters: read depth < 2; mapping quality < 40; Fisher strand bias > 60; and a Qual score < 100. High quality SNPs were annotated using snpEff (v4.3T) using the *C. auris* reference genome, B8441 (Cand_auris_B8441_V2). Genome-wide SNV-based phylogenetic analysis of the progenitor *C. auris* clinical isolates was conducted using the maximum likelihood method in RAxMLGUI (v2.0) using the variants generated above and included a total of 181229 positions. Model selection was conducted using MEGAX (v. 10.1.8) [75]. The best fitting model was the general time-reversible (GTR) model + gamma distribution. Internode support was tested using a bootstrap analysis of 500 replicates.

### Polymerase chain reaction (PCR)

All primer sequences used in both PCR and Sanger sequencing are listed in Table S6. PCR amplifications were performed with Taq DNA Polymerase (GenScript) according to the manufacturer’s instructions.

### Variant detection

*De novo* variant detection for the evolution populations after three passages in each condition was conducted and VCFs were generated using realigned, sorted BAM files with Mutect algorithm within the Genome Analysis Tool Kit (v3.7-0-gcfedb67). Variants were annotated using SnpEff (v SnpEff 4.3K, build 2017-03-29) with the *Candida auris* reference genome and gene feature file (B8441, GCA_002759435.2_Cand_auris_B8441_V2). Parental variants present in the progenitor isolates were removed using VCF-VCF intersect (https://github.com/galaxyproject/tools-iuc/tree/master/tool_collections/vcflib/vcfvcfintersect) with a window size of 0 bp. *De novo* variant detection for whole-genome sequencing of single colonies from the evolution populations was conducted with Mutect2 from the Genome Analysis Tool Kit (v4.1.2) where the progenitor strain was input as the “normal” sample and the evolved sample was the “tumor”. Low-quality variant calls were filtered with FilterMutectCalls using default parameters, and moderate and high impact variants were annotated using SnpEff. Mutect2 was also used for comparison of clade IV strains to identify alleles associated with high tolerance and high mutation rate. For the tolerance phenotype, I-P was used as the “normal” sample and H-P and Q-P were input as the “tumor” samples. For the mutator phenotype, H-P and I-P were used as “normal” samples and K-P was input as the “tumor” sample. All nonsense and missense mutations were visualized in the Integrative Genomics Viewer (IGV, v2.8.2) [37] and select variants were validated with Sanger sequencing (Table S5).

### Gene Ontology (GO) analysis

GO analysis was conducted for all terms (biological process, function, and component) using the GO Term Finder from the Candida Genome Database (CGD accessed, 10/18/2021, http://www.candidagenome.org/cgi-bin/GO/goTermFinder). Terms were considered significantly enriched if p < 0.05 with Bonferroni correction.

### Visualization of aneuploid chromosomes

Aneuploid scaffolds were visualized using the Yeast Mapping Analysis Pipeline (YMAP v 1.0) [36]. Aligned .bam files were uploaded to YMAP and read depth was plotted as a function of chromosome position using the built-in B8441 clade I reference genome. Read depth was corrected for both GC content as well as chromosome-end bias.

### Fluctuation Analysis

Fluctuation analysis of *URA3* mutation rates was performed similar to prior studies [76] using the method of the median [77]. Briefly, strains were streaked for single colonies and grown on YPAD for 2 days at 30°C. Per strain, 8 independent colonies were inoculated into 1 mL liquid YPAD and grown overnight at 30°C. Cultures were diluted with PBS and spot plated onto nonselective YPAD for total cell counts and selective media (5-FOA for *URA3* loss). YPAD plates were incubated at room temperature for 3 days and 5-FOA plates were incubated at 30°C for 3 days. Colony counts were used to calculate the rate of marker *URA3* loss per cell division.

### Data availability

The sequencing datasets generated during this study are available in the Sequence Read Archive repository under projects PRJNA635156 and PRJNA635167.

## Acknowledgements

We thank Selmecki lab members Audrey Hilk, Dalton Piotter and Liban Mohamed for technical assistance and Petra Vande Zande for critical reading of the manuscript. This work was supported by Gustavus Adolphus College Sabbatical Leave funding to L.S.B., the University of Minnesota UMR Fellowship with the Bioinformatics and Computational Biology program to N.S., the National Institutes of Health (R01AI143689) and Burroughs Wellcome Fund Investigator in the Pathogenesis of Infectious Diseases Award (#1020388) to A.S.

